# AnnoDUF: A Web-Based Tool for Annotating Functions of Proteins having Domains of Unknown Function (DUFs)

**DOI:** 10.1101/2024.06.05.597330

**Authors:** Aman Vishwakarma, Namrata Padmashali, Saravanamuthu Thiyagarajan

**Affiliations:** Institute of Bioinformatics and Applied Biotechnology (IBAB), Electronic City Phase 1, Bengaluru KA 560100, India

**Keywords:** Bioinformatics Tool, DUF – domains of unknown functions, Annotations, Web based tool

## Abstract

The rapid expansion of biological sequence databases due to high-throughput genomic and proteomic sequencing methods has left a considerable number of identified protein sequences with unclear or incomplete functional annotations. DUFs are protein domains that lack functional annotations but are present in numerous proteins. To address the challenge of finding functional annotations for DUFs, we have developed a computational method, which efficiently identifies and annotates these enigmatic protein domains by utilizing PSI-BLAST and data mining techniques. Our pipeline identifies putative potential functionalities of DUFs, thereby decreasing the gap between known sequences and functions. The tool can also take user input sequences to annotate. We executed our pipeline on 4,775 unique DUF sequences obtained from Pfam, resulting in putative annotations for 1,971 of these. These annotations were subsequently incorporated into a comprehensive database and interfaced with a web-based server named ‘AnnoDUF’. AnnoDUF is freely accessible to both academic and industrial users, via World Wide Web at the link http://bts.ibab.ac.in/annoduf.php.

## 1. INTRODUCTION

The advancements in high-throughput genomic and proteomic sequencing methods have led to a substantial increase in the scale of biological sequence databases. A significant portion of the potential proteins from these studies still lack clear or complete information regarding their functional annotations^1^. Despite significant advancements and collaborative initiatives aimed at elucidating structures and functions of proteins with unknown roles^2^ the disparity between the number of known sequences and those with known functions continues to widen. In many cases, grouping protein sequences into families based on similarities in sequence, structure, or function can assist in annotating their functions effectively. This process facilitates the identification of domain components, functional motifs, and conserved residues in terms of both structure and function and provides insights into species and sequence specific variations.

InterPro is a bioinformatics resource that through the integration of predictive models, or signatures, from multiple collaborating member databases^3^ facilitates in classification of proteins into families, highlighting functional motifs and their cellular localization. Each of the member databases utilizes a variety of distinct methods to classify proteins, with their focus on aspects such as defining protein domains from structural information or identifying full-length protein families based on shared functions. Additionally, InterPro strives to identify instances where different entries from the member databases correspond to the same entity. This meticulous approach has greatly contributed to the accuracy and reliability of protein classification and functional prediction within the InterPro database.

One of the significant contributors to the InterPro consortium is the Pfam database, widely utilized for protein family and domain analysis. Currently, Pfam has been completely integrated into Interpro and is not a stand-alone database. Pfam entries were annotated with functional information from relevant literature, wherever such information was available, through a manual process^4^. The continuous influx of new proteins has led to an increasing number of uncharacterized proteins, particularly within the category known as Domains of Unknown Function (DUFs). In the Pfam (v35.0) database, there were a total of 19,632 families, and among these, approximately 25% (Supplementary Figure S1) (4,775 out of 19,632) were categorized as DUF families, which encompassed both Domains of Unknown Function (DUF) and Uncharacterized Protein Families (UPF). Henceforth, these two families will be collectively referred to as DUF families in this article.

DUFs were first identified in the late 1990s as an increasing number of protein sequences were being discovered through genome sequencing projects. Chris Ponting pioneered the DUF naming by adding DUF1 and DUF2 to the SMART database^5,6^. It was found that a significant proportion of these sequences contained regions with no known homology to any protein with a known function. As a result, a large number of protein sequences were annotated as hypothetical or unknown. To provide a more standardized and informative annotation, DUFs were introduced to categorize these unknown protein domains. Despite their unknown function, DUFs have been found in a variety of proteins performing many important biological processes. For example, DUFs have been implicated in DNA binding, protein-protein interactions, and enzymatic activities such as hydrolase and kinase activity^7^. Furthermore, DUFs have been found to be involved in a variety of disease states, such as cancer, neurodegenerative disorders, and viral infections. As more protein sequences are discovered through advances in genome sequencing, the number of identified DUFs continues to grow (Supplementary Figure S2). It has been becoming increasingly necessary to understand these DUF proteins.

In this regard, we report here, a computational method that identifies annotations of homologues of protein with DUFs and has been developed and hosted as a web-based tool ‘AnnoDUF’. This web-based tool leverages the strengths of psi-blast and data mining techniques to annotate domains of unknown protein function (DUFs) effectively and sheds light on the elusive functionalities of these protein domains. Out of 4775 DUFs, 1971 have been given a potential annotation through our pipeline. These annotations have been integrated into a database and are made accessible to users via this tool. AnnoDUF is freely accessible to both academic and industrial users, ensuring widespread utilization of this valuable resource at http://bts.ibab.ac.in/annoduf.php. This tool can help in our understanding of the roles these domains play in protein structure and function.

## 2. MATERIALS AND METHODS

Overall pipeline is given schematically in Figure 1.

**FIGURE 1.**
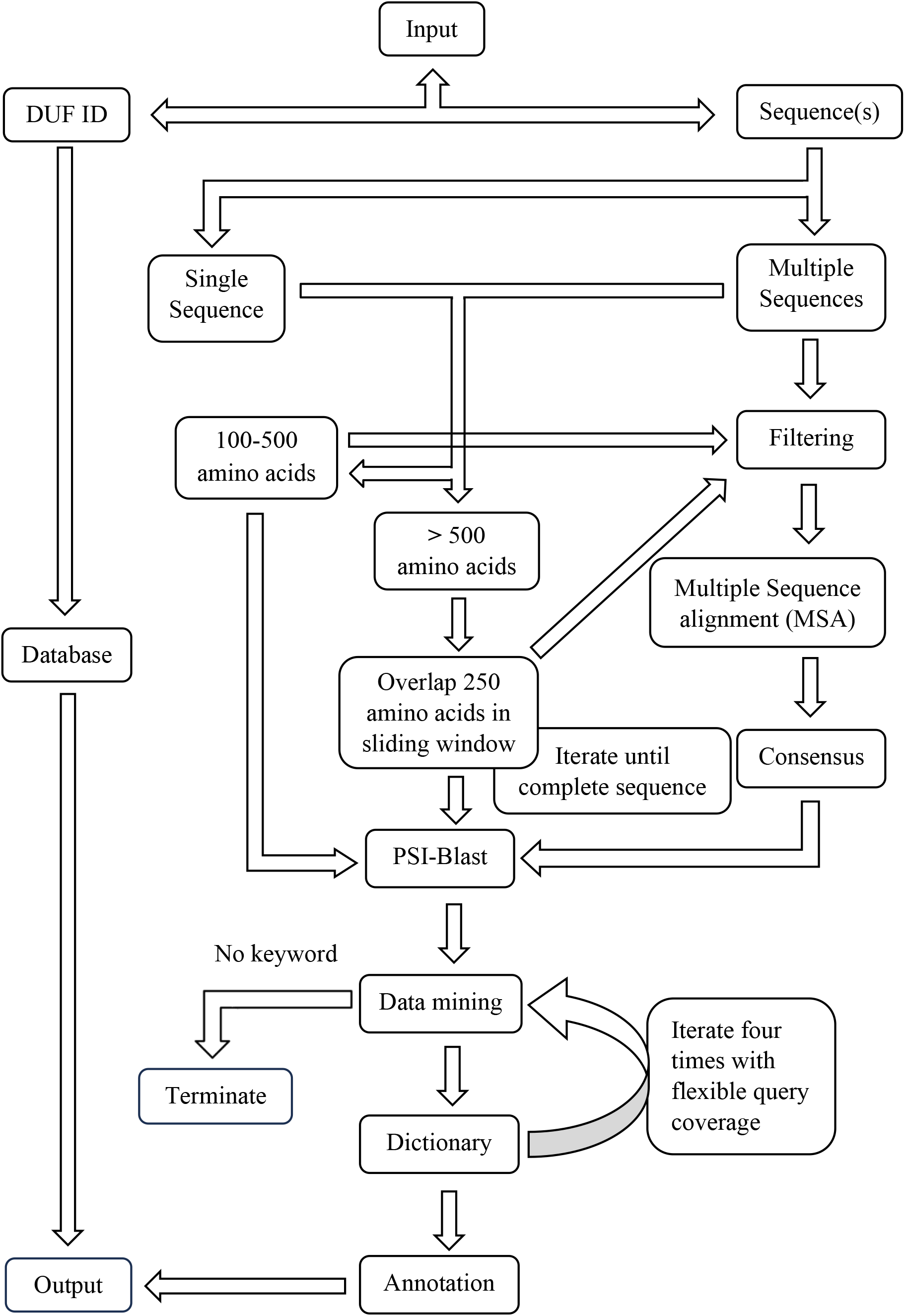
A schematic workflow of the AnnoDUF web server

### 2.1.1 Obtaining sequence data

Sequences of domains of unknown functions (DUFs) were downloaded from *Pfam* by web-scrapping using tools developed in-house or by direct download from their servers. In-house Python scripts were developed that deployed the Python modules *BeautifulSoup* (https://pypi.org/project/beautifulsoup4/), *Requests* or *Scrapy* (https://pypi.org/project/scapy/) to parse the HTML page script of the Pfam website and download the appropriate files. Then it navigates to the DUF directory using the appropriate links to extract the download links for each DUF file. Once the download links have been extracted, the script used *Requests* module again along with *Scrapy* to download each sequence file individually. The script iterates through the list of download links and uses the *get()* method to download each file. The downloaded files were saved locally on the user’s computer.

#### 2.1.2 Finding distant homologues

To include more divergent sequences or distant homologues, the query coverage range is relaxed in steps. Specifically, the query coverage range is relaxed to 70-80%, 60-70%, 50-60%, and 40-50% respectively. Each time the query coverage range is relaxed, the resulting hits were analysed to determine if there were any variations in both the quality and quantity of the results obtained. By gradually relaxing the query coverage range, additional hits were obtained that were previously excluded due to the stringent criteria.

### 2.2 Pipeline construction

A Python script has been developed to perform the annotation of DUFs, which performs the following: sequence filtering, multiple sequence alignment (MSA), deriving a consensus pattern, psi-blast, data-mining, and returning the annotation.

#### 2.2.1 Filtering the sequences

Sequence lengths of each downloaded sequence for a given DUF was calculated and the median was determined. Sequences of a given family whose length is ± 20 % of the median length was considered for further analysis. In case the resulting number of sequences is greater than 40, a random selection of 40 sequences were chosen from the filtered sequences for further processing.

#### 2.2.2 Multiple sequence alignment & Consensus sequence

Multiple sequence alignment (MSA) was performed on the filtered sequence sets, using the offline version of the *Clustal Omega* tool^8^. The choice of the scoring matrix (BLOSUM62), gap penalty scheme (affine gap penalty with a gap opening penalty of 10 and gap extension penalty of 0.1) etc., were left to the default values. The consensus sequence of the given multiple sequence alignment was generated using the EMBOSS *cons* tool^9^.

#### 2.2.3 PSI-BLAST

The consensus sequence was used as input to PSI-BLAST (offline version - 2.12.0)^10^ search against the non-redundant (*nr*) database^11^ for four iterations. Appropriate flags were used in the PSI-BLAST command to include specific output parameters, such as the query sequence ID, subject sequence accession ID, query coverage, percentage identity, alignment length, subject length, all subject titles, query start position, query end position, subject start position, subject end position, e-value, and bitscore in the output. Hits with a query coverage greater than 80 % with an identity of more than 30 % and a target coverage of 80 - 120 % were filtered and taken for data mining.

#### 2.2.4 Data-mining on PSI-BLAST result

To extract relevant information from the output, an in-house developed script was used for data mining. Common and redundant dictionary words, species names etc., were first eliminated, followed by a tabulation of frequency of each word within the list of potential hits. This enables the identification of the most frequently occurring words, which are likely to hold significant relevance for annotation. These frequently occurring words are then compiled into a word dictionary, which includes the potential hits from the list along with their respective frequencies of occurrence. The word dictionary was further refined by deleting words with a frequency less than 5, as well as words that were not required for annotation, such as “uncharacterized”, “partial”, “unknown”, and “DUF.” Additionally, alphanumeric words, such as IDs of hypothetical or uncharacterized proteins, were also deleted from the dictionary. Words containing any numeric symbols were also removed, as they are likely to represent IDs. This iterative refinement process was continued until all undesired words have been effectively eliminated from the dictionary. Once this refinement was complete, the potential hits were saved in a text file, facilitating further analysis.

#### 2.2.5 Annotations

After creating the refined word dictionary from the PSI-BLAST output, hit words in the dictionary were used as input keywords for searches in the UniProt^12^ and KEGG databases^13^ using in-house Python scripts. Relevant information from these databases were retrieved, thereby providing putative functional annotations for the target protein domain or input sequence.

### 2.3 Annotating longer sequences

The entire pipeline was executed, for families with sequences with lengths around 100-500 amino acids. For families with sequence lengths greater than 500 amino acids, the sizes were limited at a max of 500 aa. The remaining part of the sequences were included in a subsequent search, having a maximum overlap of up to 250 aa from the previous sliding window.

### 2.4 Web Interface

The pipeline has been integrated into an Apache webserver v 2.4.52 hosted on a computer running Ubuntu 22. The tool was developed using a combination of PHP (v8.1.2) for front end handling, MariaDB (v10.6.11) for the database, and Python (v3.10.6) for back-end. The interactive user interface of AnnoDUF has been designed using a combination of front-end web technologies, including XHTML 1.0 for structuring the content, CSS for styling and layout, and JavaScript to enhance interactivity. For all DUFs with or without successful annotations, the results have been integrated into a database. Thus, querying with DUF id will instantly return the results. Either the stored results or an indication that ‘no putative annotation is available’ will be returned. For sequence-based inputs, the pipeline will be executed completely and the results will be returned on the browser page. For sequence-based searches, the protein family sequence(s) can be input either through the provided text area or by uploading a file via the designated file selection box along with a job name.

The analysis begins by checking the number of sequences input protein family. In case where the input has a single sequence, the analysis proceeds without undertaking sequence filtering, multiple sequence alignment, or consensus pattern determination. However, if the family comprises more than one sequence, the analysis follows the standard methodology (Figure 1) to provide putative annotation.

### 2.6 Cross validations

For cross validating the results, a few DUFs were taken up for cross checking with other annotation tools like i) conserved domain database (CDD - https://www.ncbi.nlm.nih.gov/cdd/) ii) Pfam (currently integrated into InterPro) and iii) Annotator (https://annotator.bii.a-star.edu.sg).

## RESULTS

### 3.1 Sequence retrieval

The comprehensive collection of 4775 sequences having DUF (Domain of Unknown Function) available in the *Pfam* database, were downloaded by using an in-house developed python script. A few DUF entries did not have any candidate sequences and hence were not included in our annotation studies. Sequences for each of the DUFs were stored in separate folders for ease of calculations. The lengths of the representative protein sequences varied between 16 (DUF1752) and 39677 (DUF218) for various entries. The total number of sequences too, as collected against each DUF varied between 1 and 15955. 40 sequences per DUF were chosen randomly wherever available and were considered for further analysis. The chosen sequences had their lengths that ranged within 10 % from their respective median lengths.

### 3.2 Data Mining and Annotation

For each DUF, a multiple sequence alignment was carried out using these sets of sequences and their respective consensus sequences were deduced. This consensus sequences were considered as the representative sequences for the DUFs for downstream PSI-BLAST studies.

The PSI-BLAST iterations yielded anywhere from 3 to 4000+ homologs with an average of 500 hits per query. The titles of these PSI-BLAST results carried annotations deduced from elsewhere and were culled using data mining. This way, among the 4,775 DUF sequences taken in the current study, 1,971 were successfully annotated, providing potential functional information about them. The remaining DUF sequences that lacked putative annotations were reserved for a later run of the pipeline as new annotations get populated in the NR database of NCBI BLAST.

### 3.3 Webserver

AnnoDUF is hosted on the worldwide web and is available for all users at http://bts.ibab.ac.in/annoduf.php. The landing page is shown in Figure 2A. To initiate a search and retrieve putative annotations, the user can either enter a protein sequence in FASTA format (Figure 2B) or select the DUF ID (Figure 2C). While the former will return the results immediately, the latter option takes about 15-20 minutes to fetch results. Upon obtaining the search results, the result page presents users with an interactive table (Figure 2D) and a graphical summary (Figure 2E). The interactive summary table displays the significantly identified domains of unknown proteins that could potentially contribute to the putative annotations. The interactive graphical summary visually presents the distribution of domains within the DUFs using a pie chart (Figure 2D). Segments of the pie chart indicate the percentage of distinct domains found in the DUF sequence collection and the images also available for download with publication quality. This dynamic model enables users to rapidly comprehend the relative abundance among various domains, offering an overview of the diversity and composition of protein domains of unknown functions. Thus, users can gain valuable insights into the prevalence and significance of specific domains within the DUF dataset by interactively exploring the pie chart, assisting in the identification of potential functional clues, and facilitating further investigations into the roles of these enigmatic protein domains in biological systems. These annotated sequences were subsequently incorporated into a database for further reference and exploration. If the pipeline fails provide any annotation for a sequence-based query, an error message as in Figure 2F is returned.

**FIGURE 2.**
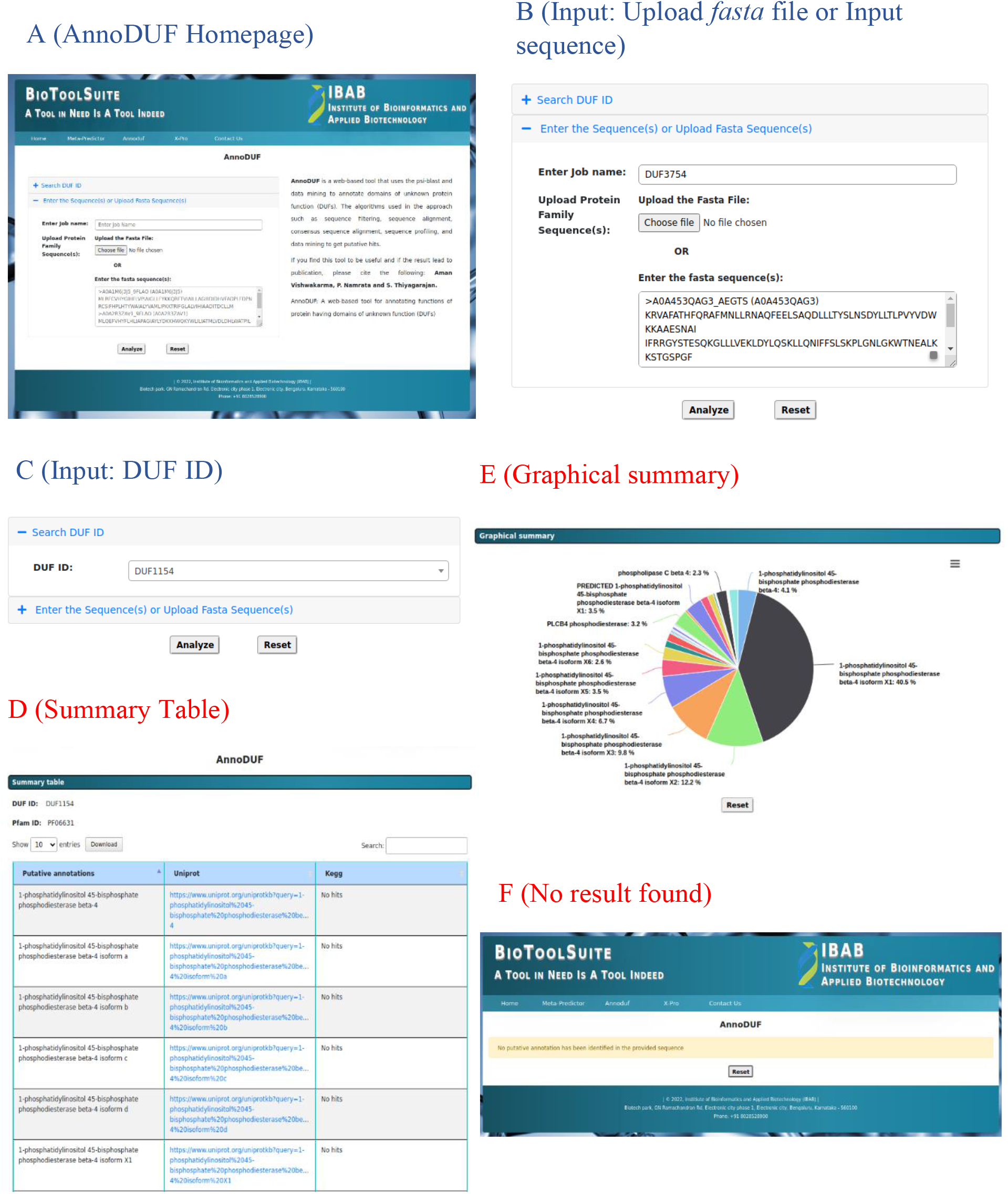
Results from AnnoDUF web server. **(A)** Landing page of the AnnoDUF webtool (**B)** Input page: protein sequence(s) can be entered in *fasta* format in a text box or a file can be uploaded. **(C)** Input option 2: A pop-up menu with an option to choose by DUF-ID. **(D)** Results of annotation are given as a summary table presenting putative annotation for the selected DUF or the uploaded sequence(s). **(E)** A Graphical summary presenting relative abundance of specific annotation obtained during the analysis. **(F)** the display message if no annotation was returned by the pipeline.

### 3.4 Assessment

To assess the accuracy of this computational method for domain annotation, the results are compared with those of other popular tools and servers. The CDD is a curated database of protein domain alignments, and it contains information on a variety of DUFs, including their sequence, structure, and function. **Table 1** lists the results of our comparison studies for the following DUFs: DUF2476, DUF1418, DUF3679, DUF2817, and DUF5614. AnnoDUF gave putative annotations to all these, whereas CDD/Pfam gave only for 3 out of 5. The Annotator tool did not yield results for any of these five randomly picked DUFs. All the annotations obtained from CDD/Pfam were consistent with AnnoDUF’s annotation whereas the other two DUFs got additional unique annotation via AnnoDUF. DUF2476 was described as proline rich protein by AnnoDUF and CDD. DUF1418 too got the annotation as member of YbJC family from AnnoDUF and CDD. DUF3679 was interpreted as some domain present in bacteria whereas AnnoDUF annotated this as YqxA family of proteins, that are involved in cell division and signalling. Likewise, DUF 2817 did not get any annotation from CDD while AnnoDUF came up with an annotation, classifying this as a member of metallopeptidases. DUF5614 is described in the CDD Database as an N-terminal domain present in C7orf25 and distantly related to PD-(D/E) nucleases. Our findings indicate that DUF5614 is present in multiple protein isoforms, including C7orf25 and UPF0415 protein C7orf25 homologs. PD-(D/E) nucleases are enzymes responsible for cleaving DNA (https://www.ebi.ac.uk/interpro/entry/InterPro/IPR011604/). A similar description was found in Pfam as well. While the exact function of DUF5614 remains unknown, it is plausible that it is associated with DNA cleavage or a related molecular process.

**Table 1:** Comparison of annotations from CDD and Pfam, Annotator from A*STAR and AnnoDUF made on a few random DUFs.

| S. No | DUF ID | CDD/Pfam | Annotator (A*STAR) | AnnoDUF |
| --- | --- | --- | --- | --- |
| 1 | DUF2476 | Proline-rich | - | Proline-rich motifs (Proline-rich 23A, Proline-rich 23B, Proline-rich 23C) |
| 2 | DUF1418 | YbjC family | - | YbjC family<br>Membrane protein |
| 3 | DUF3679 | domain present in bacteria | - | YqxA family of proteins |
| 4 | DUF2817 | - | - | M14 family of metalloproteases |
| 5 | DUF5614 | N-terminal domain present in C7orf25, PD-(D/E) nucleases | - | C7orf25, C7orf25 isoforms, UPF0415 protein C7orf25 homologs |

## DISCUSSION

The development of computational methods for domain annotation is becoming increasingly important as the number of available protein sequences continues to grow rapidly. Traditional methods of annotations, such as experimental structure determination or enzyme assays, are too slow and expensive to keep up with the pace of sequence discovery. While it may not be possible to experimentally annotate all known protein sequences, which stand at about 250 million sequences in the UNIPROT database, it may be possible to inherit the annotations of the homologous proteins, provided there is significant sequence similarity among the orthologues. By leveraging such available sequences, we can delve deeper into the mysteries of these protein domains and contribute to advancing our understanding of their roles in biological systems.

The set of 4775 sequences downloaded from Pfam encapsulates a diverse range of protein domains with unknown functions, presenting opportunities for studying their structural, functional, and evolutionary characteristics. PSI-BLAST being an authentic tool to identify distant homologues, helps in finding already annotated counterparts of any new protein sequence. The process involves filtering a large set of hits-based query coverage and percentage identify of the hits. However, the criteria are quite stringent and there may be some potentially useful annotations that are being excluded. By gradually relaxing the query coverage range, additional hits that were previously excluded due to the stringent criteria were obtained. This iterative approach results in a more comprehensive set of annotations and increased the changes of gaining valuable insights annotating unknown proteins.

The titles associated with hit sequences carry known information of the proteins. By comparing tens of or hundreds of such titles of homologues, and the keywords contained in their potential information for annotations are derived using data mining. AnnoDUF has been designed based using this data and successfully implemented as a web-server available publicly over the world wide web. AnnoDUF pipeline was executed for the total downloaded sequence and it yielded annotations for 1971 domains, which stands for more than 40 % of the unknown domains.

The development of DUF annotation tools is an important step in understanding the function of proteins. DUFs make up a significant portion of the total proteome known, and though they are all known to be involved in a variety of cellular processes. Thus, AnnoDUF stands out to be a handy tool in characterizing novel proteins, as and when they are discovered. Knowing the function of DUFs and any newly identified sequences, can help in a better understanding of the biology as well as in drug discovery.

## ACKNOWLEDGEMENTS

Authors acknowledge the institutional grant to IBAB from the Department of Electronics, IT, BT and S&T, Government of Karnataka.

## Supporting Materials

**FIGURE S1 |** The plot shows the growth of all families added to Pfam.

**FIGURE S2 |** The plot shows the growth in percentage of DUFs added to Pfam.

