## Supplementary Material for "AnnoDUF: A Web-Based Tool for Annotating Functions of Proteins having Domains of Unknown Function (DUFs)"

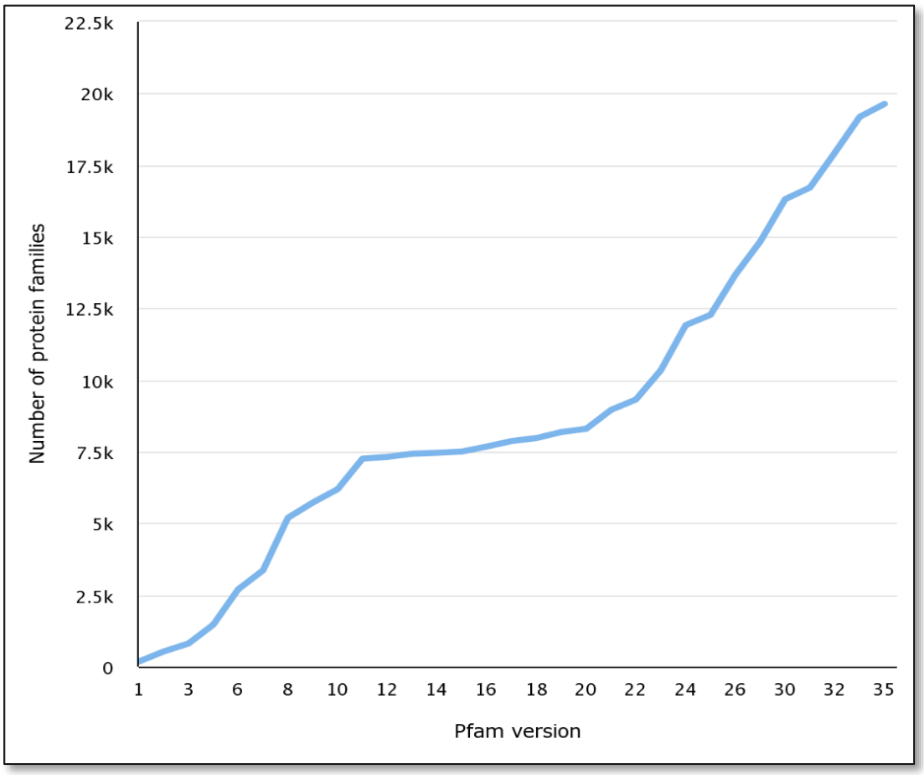


**FIGURE 1** | A graph showing the growth of all families added to Pfam.

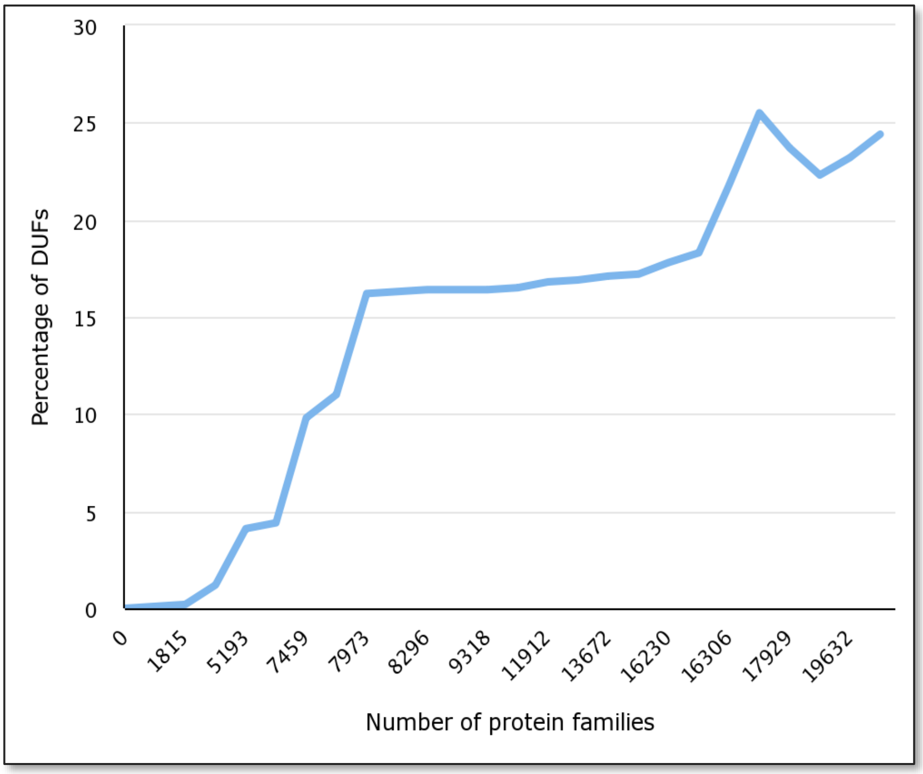


**FIGURE 2** | A graph showing the growth of all families added to Pfam.
